# Simultaneous single-channel multiplex and quantification of carbapenem-resistant genes using multidimensional standard curves

**DOI:** 10.1101/409912

**Authors:** Jesus Rodriguez-Manzano, Ahmad Moniri, Kenny Malpartida-Cardenas, Jyothsna Dronavalli, Frances Davies, Alison Holmes, Pantelis Georgiou

## Abstract

Multiplexing and absolute quantification of nucleic acids, both have, in their own right, significant and extensive use in biomedical related fields, especially in point-of-care applications. Currently, the ability to detect several nucleic acid targets in a single-reaction scales linearly with the number of targets; an expensive and time-consuming feat. Here, we propose a new methodology based on multidimensional standard curves that extends the use of real-time PCR data obtained by common qPCR instruments. By applying this novel methodology, we achieve simultaneous single-channel multiplexing and enhanced quantification of multiple targets using only real-time amplification data. This is obtained without the need of fluorescent probes, agarose gels, melting curves or sequencing analysis. Given the importance and demand for tackling challenges in antimicrobial resistance, the proposed method is applied to the four most prominent carbapenem-resistant genes: *bla*_OXA-48_, *bla*_NDM_, *bla*_VIM_ and *bla*_KPC_, which account for 97% of the UK’s reported carbapenemase-producing Enterobacteriaceae.

---

INTRODUCTION

This work builds on the framework proposed by Moniri *et al.* 2018 (1), and for the first time, demonstrates multiplex qPCR and absolute quantification employing only a single-fluorescence channel without post-PCR analysis that can be used with any PCR instrument. This is achieved by mapping real-time amplification data to a multi-dimensional space and applying novel data analytics. The proposed method is validated for the rapid screening of the *β*-lactamase genes *bla*_OXA-48_, *bla*_NDM_, *bla*_VIM_ and *bla*_KPC_ using bacterial isolates from clinical samples in a single-reaction without fluorescent probes, agarose gels, melting curves or sequencing analysis. Table 1 summarises the breakdown of confirmed carbapenemase-producing Enterobacteriaceae (CPE) cases in the UK from 2003 to 2015. The chosen drug resistant genes in this study cover over 97% of the total reported cases. Diagnostic instruments that incorporate our methodology will greatly expand the applicability of emerging molecular technologies (3, 4), including point-of-care settings.

**Table 1.** Laboratory confirmed cases of carbapenemase-producing Enterobacteriaceae from UK laboratories (2003-2015).

| Carbapenemases | *bla*<sub>OXA-48</sub> | *bla*<sub>NDM</sub> | *bla*<sub>VIM</sub> | *bla*<sub>KPC</sub> | Others |
| --- | --- | --- | --- | --- | --- |
| **Total cases** | 1325 | 1129 | 491 | 3260 | 189 |
| **Percentage (%)** | 20.73 | 17.67 | 7.68 | 51.01 | 2.96 |

Data obtained from the Public Health England's Antimicrobial resistance and healthcare associated infections reference unit (2).

Invasive infections with carbapenemase-producing strains are associated with high mortality rates (up to 40 - 50%) and represent a major public health concern worldwide (5, 6). Rapid and accurate screening for carriage of CPE is essential for successful infection prevention and control strategies as well as bed management (7, 8). However, routine laboratory detection of CPE based on carbapenem susceptibility is challenging (9): i) culture-based methods have limited sensitivity and long turnaround time; (ii) nucleic acid amplification techniques (NAATs) are often too expensive and require sophisticated equipment to be used as a screening tool in healthcare systems; and (iii) multiplexed NAATs have not been able to meet the demand for high-level multiplexing using available technologies.

There is an unmet clinical need for new molecular tools that can be successfully adopted within existing healthcare settings. The proposed method allows existing technologies to benefit from the advantages of multiplex PCR whilst reducing the complexity of CPE screening; resulting in a time and cost effective solution.

The aforementioned solution is enabled through changing the fundamental approach to current data analytic techniques for the quantification of nucleic acids from unidimensional to multidimensional. Figure 1 compares both the conventional approach (A) versus the proposed method (B). In the first, conventional standard curves are constructed but multiple targets cannot be differentiated and quantified without post-PCR processing. In contrast, the proposed method constructs multidimiensional standard curves (MSCs) extracting information from the amplification curves allowing for simultaneous quantification and multiplexing.

**Figure 1.**
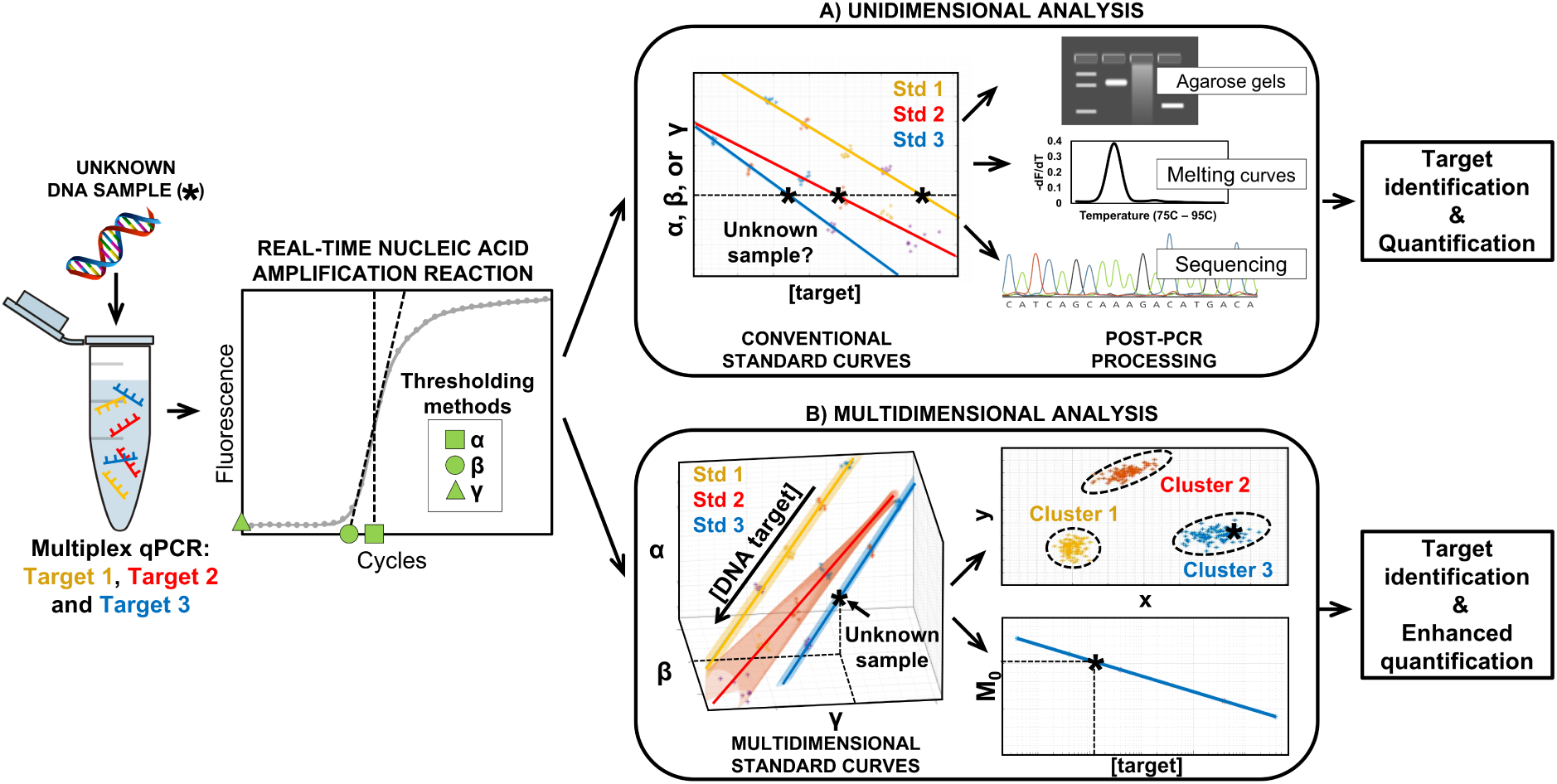
Illustration of experimental workflow for single-channel multiplex quantitative PCR using unidimensional and multidimensional approach. An unknown DNA sample is amplified by multiplex qPCR for targets 1, 2 and 3. Features such as *α*, *β* and *γ* are extracted from the amplification curve. It is important to stress that any number of targets and features could have been chosen. (A) Unidimensional analysis. Three conventional standard curves are generated through serial dilution of the known targets using a single feature. Given it is not possible to identify the target based on these standard curves, post-PCR analysis are required for target identification and quantification. (B) Multidimensional analysis. Three multidimensional standard curves are constructed through serial dilution of the known targets using multiple features. The unknown samples can be confidently classified through the use of clustering techniques and enhanced quantification can be achieved by combining all the features into a unified feature called *M*_0_.

Here, we show that the described methodology can be successfully applied to CPE screening. This provides a proof-of-concept that several nucleic acid targets can be multiplexed in a single channel using only real-time amplification data. The authors invite researchers to explore other targets and amplification chemistries in order to expand the capabilities of current state-of-the art technologies.

## MATERIALS AND METHODS

### Primers and amplification reaction conditions

All oligonucleotides used in this study were synthesised by IDT (Integrated DNA Technologies, Germany) with no additional purification. Primer names and sequences were previously reported by Monteiro *et al.* 2012 (10) (Table 2). Each amplification reaction was performed in 5 *µL* of final volume with 2.5 *µL* FastStart Essential DNA Green Master; 2 *×* concentrated (Roche Diagnostics, Germany), 1 *µL* PCR Grade water, 0.5 *µL* of 10 *×* multiplex PCR primer mixture containing the four primer sets (5 *µM* each primer) and 1 *µL* of different concentrations of synthetic DNA or bacterial genomic DNA. PCR amplifications consisted of 10 min at 95*°C* followed by 45 cycles at 95*°C* for 20 sec, 68*°C* for 45 sec and 72*°C* for 30 sec. For validation of the specificity of the products melting curve analysis was performed. One melting cycle was performed at 95*°C* for 10 sec, 65*°C* for 60 sec and 97*°C* for 1 sec (continuous reading from 65*°C* to 97*°C*). Each experimental condition was run 5 to 8 times loading the reactions into LightCycler 480 Multiwell Plates 96 (Roche Diagnostics, Germany) utilising a LightCycler 96 Real-Time PCR System (Roche Diagnostics, Germany). The concentrations of all DNA solutions were determined using a a Qubit 3.0 fluorometer (Life Technologies). Appropriate controls were included in each experiment.

**Table 2.**
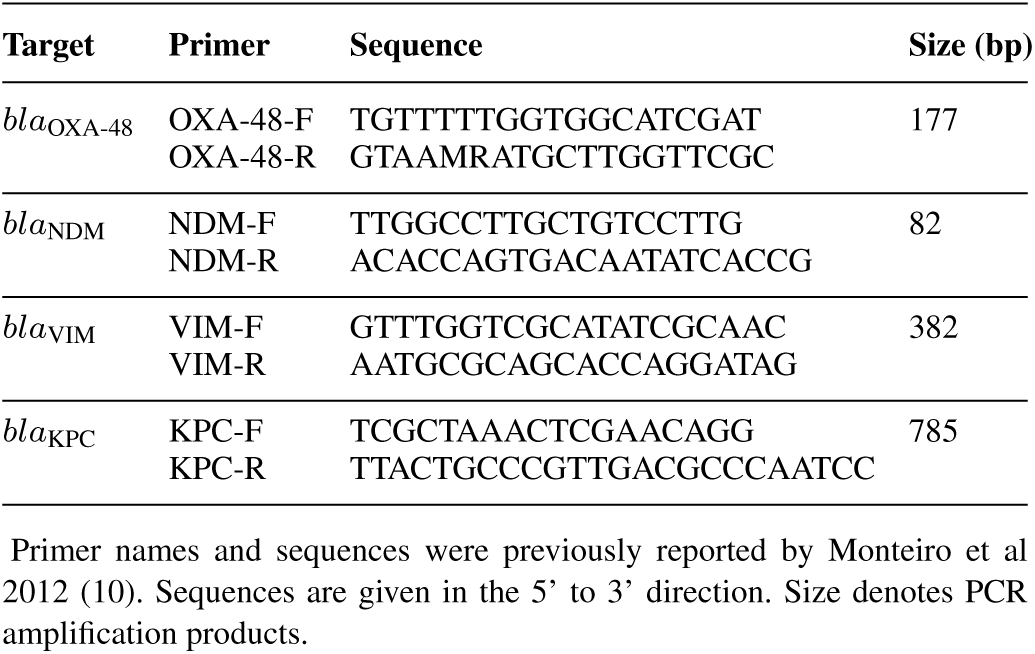
Primers used for the multiplex qPCR assay.

### Synthetic DNA and bacterial isolates

Four gBlock Gene fragments were purchased from IDT and resuspended in TE buffer to 10 *ng/µL* stock solutions (stored at −20*°C*). The synthetic templates contained the DNA sequence from *bla*_OXA-48_, *bla*_NDM_, *bla*_VIM_ and *bla*_KPC_ genes required for the multiplex qPCR assay (Supplementary Information, sheet 1). Eleven pure bacterial cultures from clinical isolates were used in this study, as described in Table 3. One loop of colonies from each pure culture was suspended in 50 *µL* digestion buffer (Tris-HCl 10 mmol/L, EDTA 1 mmol/L, pH 8.0 containing 5 *U/µL* lysozime) and incubated at 37*°C* for 30 min in a dry bath. subsequently, 0.75 *µL* proteinase K at 20 *µg/µL* (Sigma) were added, and the solution was incubated at 56*°C* for 30 min. After boiling for 10 min to inactivate proteinase K, the samples were centrifuged at 10,000 *×* g for 5 min and the supernatant was transferred in a new tube and stored at −80*°C* before use. Sample 9 was generated by mixing sample 6 & 8 at equal proportions. Bacterial isolates included non-CPE producer *Klebsiella pneumoniae* and *Escherichia coli* as control strains.

**Table 3.** Bacterial isolates used in this study.

| Sample ID | Bacterial Isolate | Carbapenemase genes |
| --- | --- | --- |
| 1 | *Klebsiella pneumoniae* | *bla*<sub>OXA-48</sub> |
| 2 | *Escherichia coli* | *bla*<sub>OXA-48</sub> |
| 3 | *Citrobacter Freundii* | *bla*<sub>VIM</sub> |
| 4 | *Escherichia coli* | *bla*<sub>NDM</sub> |
| 5 | *Klebsiella pneumoniae* | *bla*<sub>OXA-48</sub> |
| 6 | *Klebsiella pneumoniae* | *bla*<sub>NDM</sub> |
| 7 | *Pseudomonas aeruginosa* | *bla*<sub>VIM</sub> |
| 8 | *Klebsiella pneumoniae* | *bla*<sub>KPC</sub> |
| 9 | *Klebsiella pneumoniae* | *bla*<sub>NDM</sub> + *bla*<sub>KPC</sub> |
| 10 | *Klebsiella pneumoniae* | non-producer |
| 11 | *Escherichia coli* | non-producer |

### Instance of framework

The data analysis for simultaneous quantification and multiplexing is achieved using the framework described in Moniri *et al.* 2018 (1). Therefore, there are multiple stages for data analysis that are briefly described and referenced in Table 4. For more details, please refer to supplementary data (methods file).

**Table 4.** Instance of framework proposed in Moniri *et al.* 2018 (1).

| Data analysis stages | Method | Ref. |
| --- | --- | --- |
| Pre-processing | Baseline correction | - |
| Curve fitting | 5-parameter sigmoid | (11) |
| Feature extraction | $C_t$, $C_y$ and $\log_{10}(F_0)$ | (12,13,14) |
| Line fitting | Method of least squares | (15) |
| Similarity measure. | Mahalanobis distance: d | (1,16) |
| Feature weights | Minimise figure of merit: Q | (1) |
| Dimensionality reduction | Principal component regression: M0 | (15) |

## RESULTS

In this study, it is shown that simultaneous enhanced quantification and multiplexing of *bla*_OXA-48_, *bla*_NDM_, *bla*_VIM_ and *bla*_KPC_ *β*-lactamase genes in bacterial isolates can be achieved through analysing the fluorescent amplification curves in qPCR by using multidimensional standard curves. This section is broken into two parts: target discrimination using multidimensional analysis and enhanced quantification. First, it is proven that single-channel multiplexing can be achieved. Once this has been established, the framework described in Moniri *et al.* (2018) (1) can be applied for robust and enhanced quantification.

### Target Discrimination using Multidimensional Analysis

Given that it is non-trivial that several targets can be multiplexed and differentiated using only fluorescent amplification data in a single channel, it is helpful to visualise an example. Figure 2 shows four amplification curves and their respective derived melting curves specific for *bla*_OXA-48_, *bla*_NDM_, *bla*_VIM_ and *bla*_KPC_ genes. The four curves have been chosen to have similar *C*_*t*_ (within 1.2 cycle). Using only this information, i.e. the conventional way of thinking, post-PCR processing such as melting curve analysis is needed to differentiate the targets. The same argument applies when solely observing *C*_*y*_ and *F*_0_. This is an expected result given that these parameters are used for quantification and were not developed for target identification.

**Figure 2.**
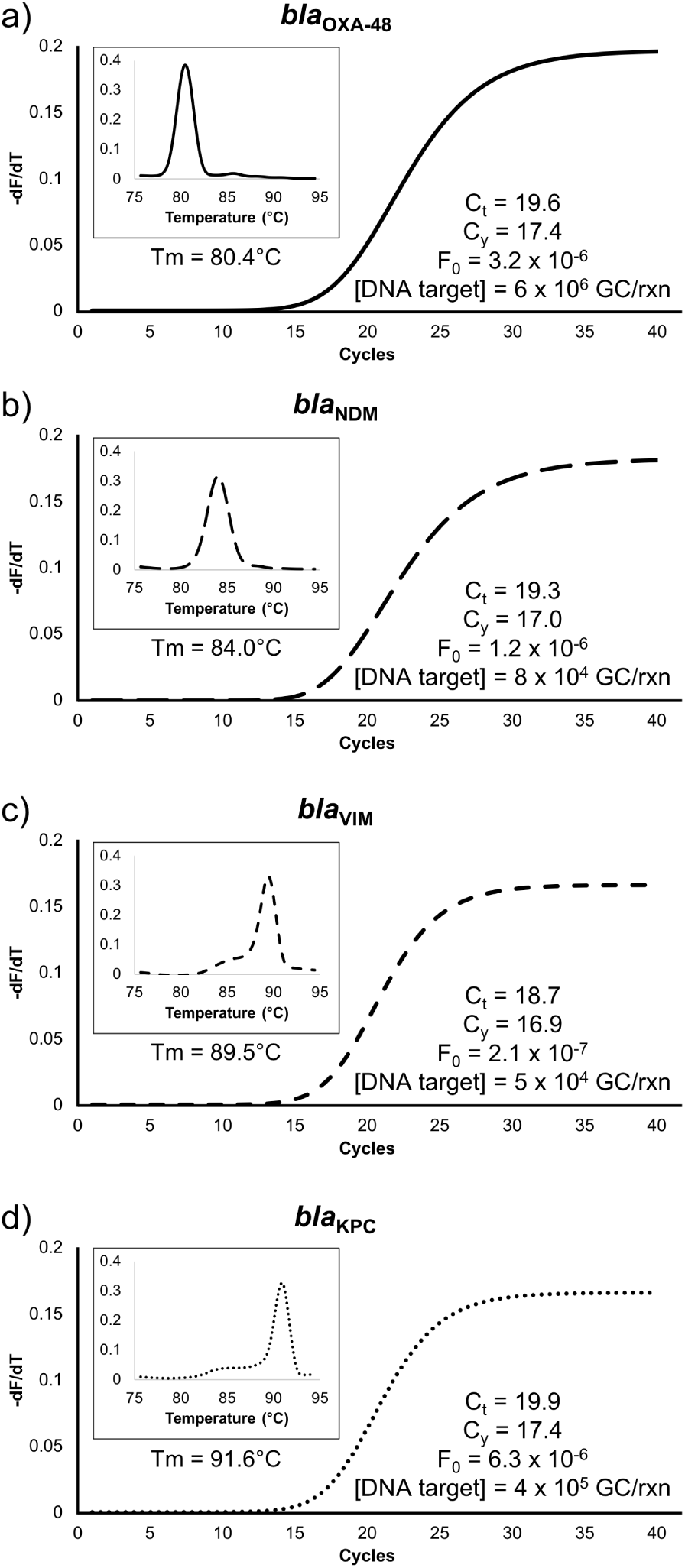
Experimentally obtained amplification and melting curves by single-channel multiplex quantitative PCR for *bla*_OXA-48_, *bla*_NDM_, *bla*_VIM_ and *bla*_KPC_ genes. Extracted features (*C*_*t*_, *C*_*y*_ and *F*_0_) are shown on the respective plots. Each plot has been generated by amplifying standard synthetic DNA containing the target of interest. All samples contained SYBR Green I dye.

Moniri *et al.* 2018 (1) shows that considering multiple features contains sufficient information gain in order to discriminate outliers from the specific target using a MSC. However, this raises the question: does the outlier lie on its own MSC? If so, can we take advantage of this property and build several multidimensional standard curves in order to discriminate multiple specific targets?

In order to explore this new concept, MSCs are constructed using a single primer mix for the four target genes using *C*_*t*_, *C*_*y*_ and *- log*_10_(*F*_0_), as shown in Figure 3. It is visually observed that the 4 standards are sufficiently distant in multidimensional space, also termed the feature space, in order to distinguish them. That is, an unknown DNA sample can be potentially classified as one of the specific targets (or an outlier) solely using the extracted features from amplification curves in a single channel.

**Figure 3.**
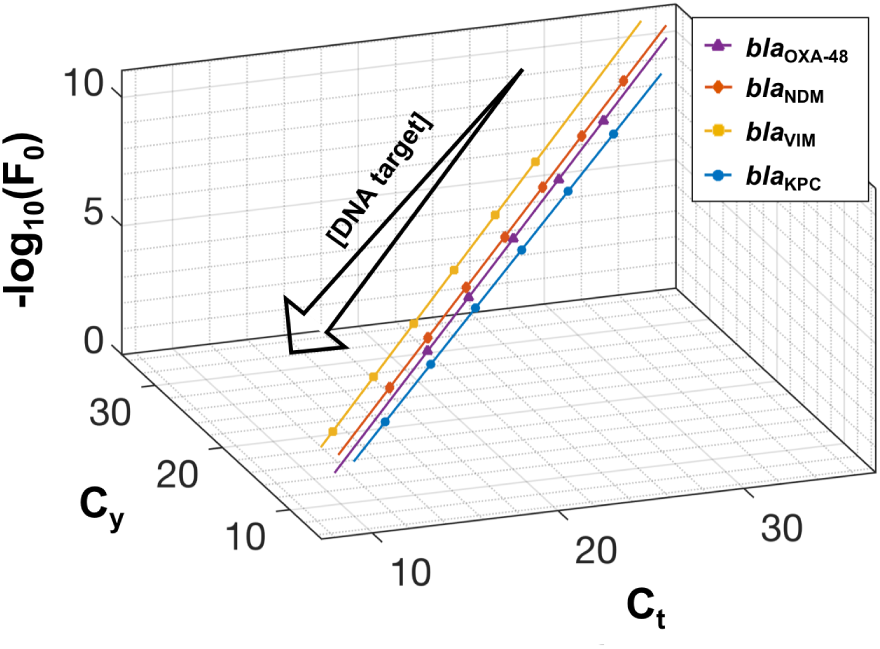
Multidimensional standard curves for detection and quantification of four carbapenemase genes: *bla*_OXA-48_ (purple line), *bla*_NDM_ (red line), *bla*_VIM_ (yellow line) and *bla*_KPC_ (blue line). They were constructed using *C*_*t*_, *C*_*y*_ and *-log*_10_(*F*_0_) features extracted from real-time amplification curves derived from amplifying 10-fold dilutions of synthetic DNA. From top right to bottom left, target concentrations range between: 6 x 10^3^ - 6 x 10^7^ for *bla*_OXA-48_; 8 x 10^0^ - 8 x 10^6^ for *bla*_NDM_; 5 x 10^2^ - 5 x 10^7^ for *bla*_VIM_; and 4 x 10^2^ - 4 x 10^7^ for *bla*_KPC_. Each concentration was repeated 5 to 8 times and the resulting average values are projected onto the standard curves. The computed features and curve-fitting parameters for each MSC is presented in supplementary data (sheet 2).

In order to demonstrate the proposed method, 11 samples (bacterial isolates) given in Table 3 were tested against the multidimensional standards. The similarity measure used to classify the unknown samples is the Mahalanobis distance using a p-value of 0.01 as the threshold to determine if the sample is an outlier. Figure 4 shows the Mahalanobis space for the four standards. This visualisation is constructed by projecting all data points onto an arbitrary hyperplane orthogonal to each standard curve, as described in Moniri *et al.* 2018 (1).

**Figure 4.**
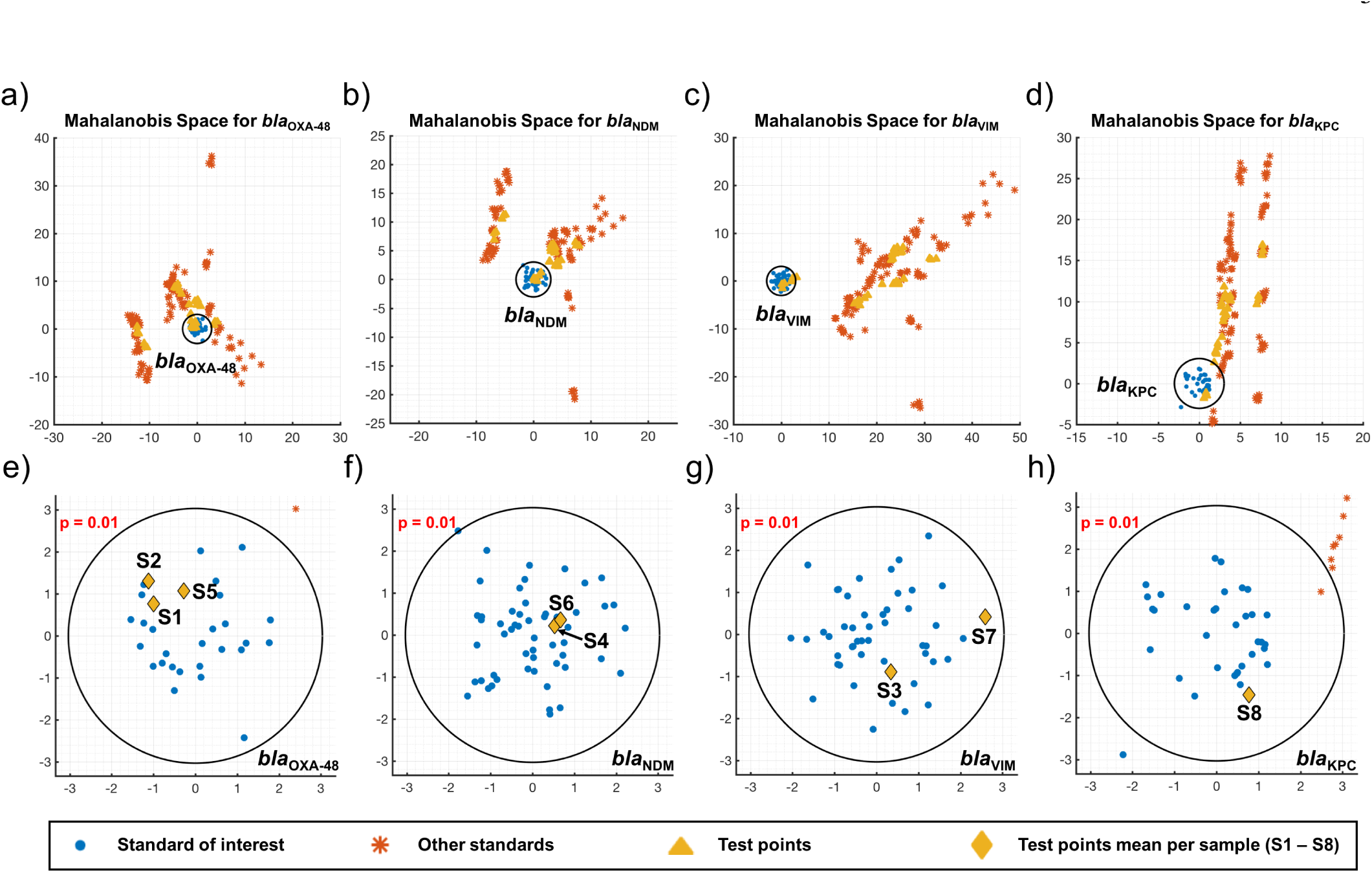
Multidimensional analysis using the feature space for clustering and classification of unknown samples. (a-d) Arbitrary hyperplanes orthogonal to each multidimensional standard curve have been used to project all the data points, including the replicates for each concentration for the four multidimensional standards (training standard points) and nine unknown samples (test points). (e-h) The previous plots are magnified to visualise the location of the samples relative to each standard of interest. All samples are classified correctly. The blue points represent the standard of interest (5 to 8 replicates per each concentration). Black circle corresponds to a p-value of 0.01. Please see supplementary data (sheet 4 and 5) for more details.

The first observation is that the training points (synthetic DNA) from each standard curve are clustered together (i.e. not considered outliers) in its respective Mahalanobis space yet they are considered outliers for other MSCs. This corroborates the fact that there is sufficient information in the 3 chosen features to distinguish the 4 standard curves. The second observation is that the mean of the test samples (bacterial isolates) which have a single resistance (samples 1-8) fall within the correct cluster of training points. Melting curve analysis was used to validate the results, as provided in supplementary data (sheet 3). The results from testing can be succinctly captured within a bar chart shown in Figure 5. However, it is important to visualise the data in order to confirm that the Mahalanobis distance is a suitable similarity measure. When the training data points in the feature space are approximately normally distributed, then the distribution of the training data points in the Mahalanobis space is approximately circular - as seen in Figure 4.

**Figure 5.**
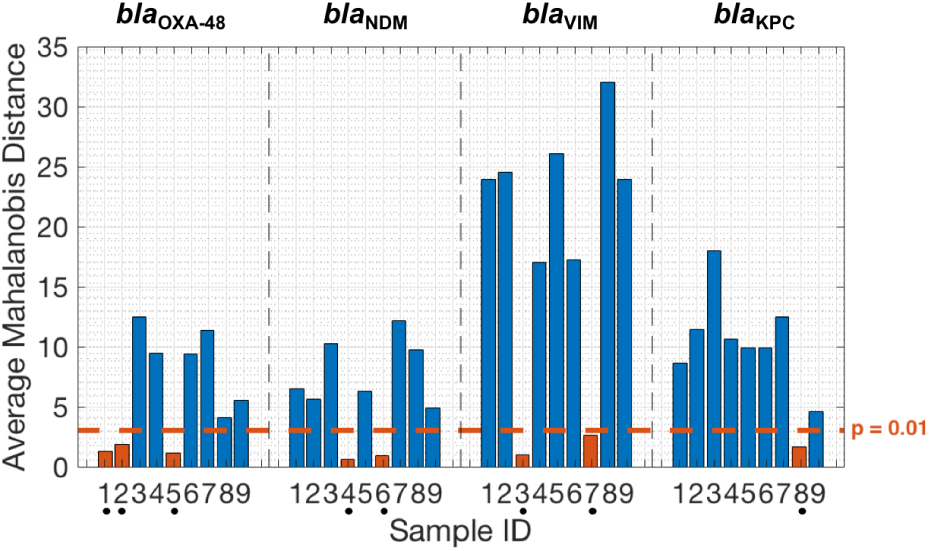
Average Mahalanobis distance from standard training points to sample test points (bacterial isolates) for target identification. Dots below sample ID indicate that the test sample is classified to the standard of interest - as indicated above the plot - with a p-value *<* 0.01.

Another interesting observation is regarding sample 9 - a target with multiple resistances. This sample, as seen in Figure 5, is considered as an outlier (p-value *<* 0.01) for all of the standards. The presence of multiple resistant genes was validated using melting curve analysis in supplementary data (sheet 3).

### Enhanced Quantification

Given that multiplexing has been established, quantification can be trivially obtained using any conventional method such as the gold standard cycle threshold, *C*_*t*_. However, as shown in Moniri *et al.* 2018 (1), enhanced quantification can be achieved using a feature, *M*_0_, that combines all of the features. This is achieved through weighting each feature by optimising an objective function and then applying a dimensionality reduction technique in order to create a quantification curve for *M*_0_. The objective function in this study is a figure of merit that combines accuracy, precision, stability and overall predictive power as described in detail in supplementary data (methods file). The weights, *α*, and quantification curves for each MSC can be found in supplementary data (sheet 6). Table 5 shows the figure of merit for the 3 chosen features (*C*_*t*_, *C*_*y*_ and *-log*_10_(*F*_0_)) and *M*_0_. The percentage improvement is also shown. It can be observed that quantification is always improved compared to the best single feature. The improvement is 65.5%, 34.4%, 54.7% and 64.9% for *bla*_OXA-48_, *bla*_NDM_, *bla*_VIM_ and *bla*_KPC_ respectively. This is expected given the nature of the multidimensional framework. However, it is interesting to observe that amongst the conventional methods, there is no single method that performs the best for *all* the targets. Thus, *M*_0_ is the most robust method in the sense that it will always be the best performing method.

**Table 5.** Figure of merit comparing conventional features with *M*_0_ for absolute quantification.

|  | *bla*<sub>OXA-48</sub> | *bla*<sub>NDM</sub> | *bla*<sub>VIM</sub> | *bla*<sub>KPC</sub> |
| --- | --- | --- | --- | --- |
| $C_t$ | **533.4** | 64.6 | **53.2** | 3851.7 |
| $C_y$ | 540.1 | **49.3** | 54.4 | **1883.1** |
| $F_0^*$ | 1985.6 | 22646.4 | 1903.7 | 40683.2 |
| $M_0$ | **183.8** | **32.3** | **24.1** | **660.9** |
| % Imp. | 65.5 | 34.4 | 54.7 | 64.9 |
| p-value | 0.04 | 0.07 | 0.01 | 0.04 |

% Imp. = Percentage improvement of $M_0$ over the next best method (both in bold). \*The figure of merit values is calculated using $-\log_{10}(F_0)$ . P-values are computed using a paired t-test (two-tailed).

## DISCUSSION

There has been a long-standing goal to meet the demand of methods for high-level multiplexing and enhanced quantification. Any advancements in this area would have a substantial positive impact on healthcare and patient outcomes. Alongside these challenges, there is a growing concern around antimicrobial resistance; the past 10 years has seen an explosion in molecular methods and instruments for rapid screening of drug resistant genes (17). Here, we propose a novel method that redefines the foundations of qPCR data analysis, allowing for simultaneous single channel multiplexing and enhanced quantification using existing technologies without the need of post-PCR processing or fluorescent probes. This method has been validated for the rapid screening of carbapenemase genes and represents the achievement of a new milestone in nucleic acid based diagnostics.

Current methods for multiplexing and quantifying involve either using fluorescent probes, melting curve analysis, agarose gels or sequencing - all of which are time-consuming and expensive processes. In the last few years there have been attempts to achieve simultaneous multiplexing and quantification in a single channel. Single channel multiplexing has been achieved without melting curve analysis by altering cycling conditions and reading fluorescence at different temperatures (18). This resulted in sufficient information gain in order to discriminate two targets. However, research has been artificially limited by a unidimensional way of thinking. Therefore, modifications in PCR conditions had to be applied, extending reaction time and potentially damaging reaction efficiency, as apposed to extracting information from existing data using more complex data analytics; incurring no extra cost or assay time.

By performing multidimensional analysis, we only require: to build multiple multidimensional standard curves (a one-time procedure); and data processing (such as multi-feature extraction) which is negligible given the power of computers today. In addition, given the nature of the multidimensional framework, absolute quantification is enhanced through the use of *M*_0_ by optimising a figure of merit combining accuracy, precision, stability and overall predictive power. The authors invite researchers in this area to adopt *M*_0_ for absolute quantification as it guarantees improved performance by combining the benefits of all the features it is derived from. This property results in *M*_0_ offering the most robust method of quantification as it provides the best quantification performance across targets.

Given the novelty of this work, there are many future directions and questions that can be addressed. In this paper we have applied the proposed method to the rapid screening of the four most prominent carbapenemase genes. In future studies, it would be interesting to explore: other targets in order to develop new multiplexing panels associated with the most significant healthcare challenges; more targets through incorporating additional multidimensional standard curves into the feature space or using multiple fluorescent channels. It is also important to stress that the focus of this work was not on optimising the chemistry or data analytics for this specific set of targets. Thus, there is room to investigate whether the chemistry and the instance of framework can be optimised in order to maximise the separation of MSCs in the feature space for carbapenem–resistant genes.

In conclusion, this work has shown that is possible to simultaneously quantify and multiplex several targets in a single channel using only amplification data obtained from existing technologies by changing the way we think about data into higher dimensions. We hope that by sharing these ideas, researchers and practitioners can implement and advance this work in order to provide novel and affordable tools that can be easily adopted by healthcare systems.

## REFERENCES

1. Moniri, A., Rodriguez-Manzano, J., Georgiou, P. (2018) A framework for the analysis of real-time nucleic acid amplification data using novel multidimensional standard curves. Preprint at bioRxiv. https://doi.org/10.1101/379180

2. GOV. UK. (2018). Carbapenemase-producing Enterobacteriaceae: laboratory confirmed cases. [online] Available at: https://www.gov.uk/government/publications/carbapenemase-producing- enterobacteriaceae-laboratory-confirmed-cases [Accessed 1 May 2018].

3. Moser, N., Rodriguez-Manzano, J., Lande, T.S. and Georgiou, P. (2018) A Scalable ISFET Sensing and Memory Array With Sensor Auto-Calibration for On-Chip Real-Time DNA Detection. IEEE transactions on biomedical circuits and systems, 12, 390–401.

4. Moser, N., Rodriguez-Manzano, J., Yu, L.S., Kalofonou, M., Mateo, S.d., Li, X., Lande, T.S., Toumazou, C. and Georgiou, P. (2017) Live demonstration: A CMOS-based ISFET array for rapid diagnosis of the Zika virus, 2017 IEEE International Symposium on Circuits and Systems (ISCAS), pp. 1–1.

5. van Duin, D. and Doi, Y. (2017) The global epidemiology of carbapenemase-producing Enterobacteriaceae. Virulence, 8, 460–469.

6. Otter, J.A., Burgess, P., Davies, F., Mookerjee, S., Singleton, J., Gilchrist, M., Parsons, D., Brannigan, E.T., Robotham, J. and Holmes, A.H. (2017) Counting the cost of an outbreak of carbapenemase-producing Enterobacteriaceae: an economic evaluation from a hospital perspective. Clinical microbiology and infection: the official publication of the European Society of Clinical Microbiology and Infectious Diseases, 23, 188–196.

7. Otter, J.A., Doumith, M., Davies, F., Mookerjee, S., Dyakova, E., Gilchrist, M., Brannigan, E.T., Bamford, K., Galletly, T., Donaldson, H. et al. (2017) Emergence and clonal spread of colistin resistance due to multiple mutational mechanisms in carbapenemase-producing Klebsiella pneumoniae in London. Scientific reports, 7, 12711.

8. Carmeli, Y., Akova, M., Cornaglia, G., Daikos, G.L., Garau, J., Harbarth, S., Rossolini, G.M., Souli, M. and Giamarellou, H. (2010) Controlling the spread of carbapenemase-producing Gram-negatives: therapeutic approach and infection control. Clinical microbiology and infection: the official publication of the European Society of Clinical Microbiology and Infectious Diseases, 16, 102–111.

9. Viau, R., Frank, K.M., Jacobs, M.R., Wilson, B., Kaye, K., Donskey, C.J., Perez, F., Endimiani, A. and Bonomo, R.A. (2016) Intestinal Carriage of Carbapenemase-Producing Organisms: Current Status of Surveillance Methods. Clinical Microbiology Reviews, 29, 1–27.

10. Monteiro, J., Widen, R.H., Pignatari, A.C., Kubasek, C. and Silbert, S. (2012) Rapid detection of carbapenemase genes by multiplex real-time PCR. J Antimicrob Chemother, 67, 906909.

11. Spiess, A.-N., Feig, C., and Ritz, C. (2008). Highly accurate sigmoidal fitting of real-time PCR data by introducing a parameter for asymmetry. BMC Bioinformatics, 9, 221.

12. Wittwer, C.T., Herrmann, M.G., Moss, A.A. and Rasmussen, R.P. (1997) Continuous fluorescence monitoring of rapid cycle DNA amplification. Biotechniques, 22, 130138.

13. Guescini, M., Sisti, D., Rocchi, M.B., Stocchi, L. and Stocchi, V. (2008) A new real-time PCR method to overcome significant quantitative inaccuracy due to slight amplification inhibition. BMC Bioinformatics, 9, 326.

14. Rutledge, R.G. (2004) Sigmoidal curve-fitting redefines quantitative real-time PCR with the prospective of developing automated high-throughput applications. Nucleic Acids Res, 32, e178e178.

15. Friedman, J., Hastie, T. and Tibshirani, R. (2001). The elements of statistical learning. New York: Springer series in statistics, pp. 337387.

16. De Maesschalck, R., Jouan-Rimbaud, D. and Massart, D.L. (2000) The Mahalanobis distance. Chemometr Intell Lab Syst, 50, 118.

17. Osei Sekyere, J., Govinden, U. and Essack, S.Y. (2015) Review of established and innovative detection methods for carbapenemase-producing Gram-negative bacteria. Journal of applied microbiology, 119, 1219–1233.

18. Lee, Y.-J., Kim, D., Lee, K. and Chun, J.-Y. (2014) Single-channel multiplexing without melting curve analysis in real-time PCR. Scientific reports, 4, 7439.

